# Uncertainty Quantification in Variable Selection for Genetic Fine-Mapping using Bayesian Neural Networks

**DOI:** 10.1101/2022.02.23.481675

**Authors:** Wei Cheng, Sohini Ramachandran, Lorin Crawford

## Abstract

In this paper, we propose a new approach for variable selection using a collection of Bayesian neural networks with a focus on quantifying uncertainty over which variables are selected. Motivated by fine-mapping applications in statistical genetics, we refer to our framework as an “ensemble of single-effect neural networks” (ESNN) which generalizes the “sum of single-effects” regression framework by both accounting for nonlinear structure in genotypic data (e.g., dominance effects) and having the capability to model discrete phenotypes (e.g., case-control studies). Through extensive simulations, we demonstrate our method’s ability to produce calibrated posterior summaries such as credible sets and posterior inclusion probabilities, particularly for traits with genetic architectures that have significant proportions of non-additive variation driven by correlated variants. Lastly, we use real data to demonstrate that the ESNN framework improves upon the state-of-the-art for identifying true effect variables underlying various complex traits.

## 1 Introduction

Variable selection is a fundamental problem in high-dimensional statistical learning that arises in a wide range of application domains [1, 2, 3, 4]. An important benefit of incorporating sparsity when building a predictive model is that it provides interpretations on which input variables are most important in explaining variation across the output variables. Such a property is particularly desirable when the end goal of an application also includes scientific discovery. For example, the goal of many genome-wide association (GWA) studies is not just to predict the disease status or phenotypic risk of a patient but also to identify the (subsets of) single nucleotide polymorphisms (SNPs) that are statistically associated with the genetic architecture of the disease [5, 6]. This can further help with downstream clinical applications such as drug development.

While many methods for variable selection have been developed in the literature [1, 2, 3, 4, 7, 8], some significant challenges still remain. One important challenge is assessing the uncertainty in which variables should be selected when they are highly correlated [9, 3]. As an extreme case, imagine there are two variables that are completely collinear. In this context, it becomes statistically impossible to distinguish them, and many traditional regularization and shrinkage methods will arbitrarily select one SNP as being associated with the trait of the interest and disregard the other [9]. While such a strategy suffices if the goal is to build a predictive model, it becomes limiting for scientific discovery because the conclusions rely on selecting the correct subset of genetic variants for downstream investigation. Recently, Wang et al. [9] introduced the “sum of single-effects” model called SuSiE to address these issues. More specifically, SuSiE assesses the uncertainty of variables by providing “credible sets” which, in the case of our extreme example, effectively summarize that “either SNP 1 or 2 are relevant but we are unsure as to which one”. SuSiE uses an iterative Bayesian stepwise selection (IBSS) procedure where it will iteratively regresses out effect variables and feeds the corresponding residuals to the next iteration for training.

The main limitation of SuSiE is that it is a linear model and therefore does not capture nonlinear effects in data. In GWA studies, it is well known that the genetic architecture of complex traits can be driven by phenomena such as dominance and epistasis [10, 11, 12, 13]. Indeed, machine learning models are most powered in settings when large sets of training data are available and often exhibit greater predictive accuracy than linear models in applications driven by non-additive variation. In this paper, we introduce the “ensemble of single-effect neural networks” (ESNN) framework which overcomes the limitations of SuSiE while preserving the ability to assess uncertainty for variable selection. We demonstrate our approach in a simulation study and on two real GWA datasets.

## 2 The Sum of Single-Effects Regression Model

In this section, we provide background on single-effects regression (SER) and state a rigorous definition of credible sets for variable selection. The original SER model [14, 15] assumes that exactly one of *J* input variables has a non-zero coefficient. More specifically,

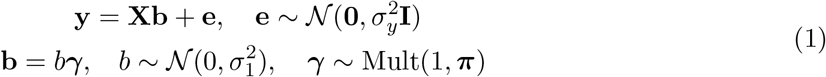

where **y** is an *N*-dimensional response vector (e.g., continuous phenotypes); **X** is an *N* × *J* design matrix (e.g., genotypes); **e** is an *N*-dimensional error term; **b** is a *J*-dimensional vector of regression coefficients; *γ* is a binary indicator that determines which regression coefficient is to be non-zero; and Mult(*m, π*) denotes the multinomial distribution with *m* samples drawn with class probability distribution *π*. For simplicity, we will consider a uniform prior such that π = (1/*J*,…,1/*J*). Note that m is set to equal to one so that the coefficient vector **b** has exactly one non-zero entry for modeling the single-effect. To estimate the statistical association of each variable, one would fit *J*-univariate models corresponding to regressing each *j*-th column **x**_*j*_ of **X** onto the response **y** and computing posterior inclusion probabilities defined as 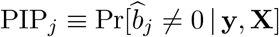.

In the context of statistical genetics, the original SER model only assumes one causal SNP. However, we know that many real-world applications, it is desired to have a method that flexibly allows for many variants to have an effect on trait architecture [3, 16]. The SuSiE framework is based on an extension of summing over *L*-multiple SER models [9]. Here, the main idea is to construct an overall effect vector **b** from multiple single-effect coefficients **b**_1_,…, **b**_*L*_ via the following

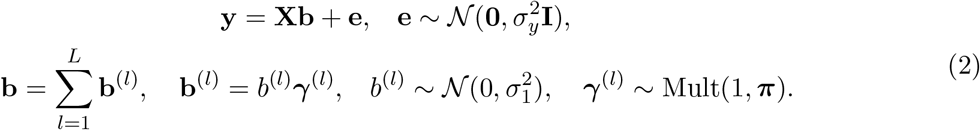

In practice, SuSiE uses an iterative Bayesian stepwise selection (IBSS) algorithm (i.e., coordinate ascent variational inference) to estimate the model parameters. More specifically, at each iteration, it fits the SER model for **b**^(*l*)^ using the residuals from the model 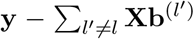. At the end of training, the SuSiE model provides *L* estimated coefficient vectors 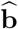 and *L* corresponding PIP vectors 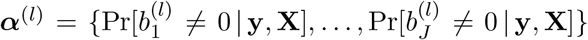. Computation of a final inclusion probability assumes that effects are independent across the *L* different models and is computed as

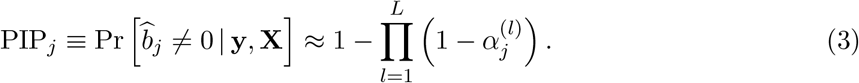

A key component of SuSiE is that it uses these PIPs to naturally construct credible sets. Effectively, a level *ρ* credible set 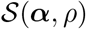 can be estimated by simply sorting variables in descending order and then including variables into the set until their cumulative probability exceeds *ρ* [9]. Below we give the rigorous definition for credible sets.

#### Definition 1

**(Wang et al. (2020) [9])** *In the context of a multiple-regression model, a level ρ credible set is defined to be a subset of variables that has probability ρ or greater of containing at least one effect variable (i.e., a variable with non-zero regression coefficient). Equivalently, the probability that all variables in the credible set have zero regression coefficients is* 1 – ρ *or less*.

The definition above yields a metric for assessing the uncertainty when conducting variable selection. A credible set will determine if a subset of collinear variables have effects on the response even when we are unclear as to which specific ones. This differs from the results produced by the conventional regularization and shrinkage methods [3, 8, 7] where the effect sizes for an arbitrarily selected subset of correlated variables will be penalized while the others are retained.

## 3 The Ensemble of Single-Effect Neural Networks

In this section, we detail the full specification of our proposed nonlinear framework for variable selection. While there exist many nonlinear models, neural networks are well known to have the ability to approximate complex systems [17, 18]. For simplicity, we will focus on multi-layer perceptrons throughout this paper; however, we also want to emphasize that the theoretical concepts we describe can also be applied broadly to other architectures (e.g., convolutional neural networks). Formally, we specify a *K*-layer probabilistic neural network as a generalized nonlinear model

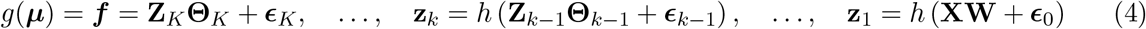

where, in expectation, the response variable is related to the input data by 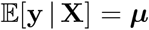; ***f*** is an *N*-dimensional latent vector to be learned; *g*(•) denotes a general cumulative link function which, for example, is set to be the identity if **y** is continuous or the logit if **y** is binary; **Z**_*k*_ denotes the matrix of nonlinear neurons from the k-th hidden layer with corresponding weight matrix **Θ**_*k*_; *ϵ_k_* are deterministic biases that are produced during the network training phase for the *k*-th hidden layer; *h*(•) is a nonlinear activation function (e.g., ReLU or tanh); and **W** is a matrix of weights for the input layer.

Similar to the SER model, the key design that leads to our ability to model single-effect is through the prior we place on the input layer weights in **W**. Let *H_k_* represent the number of neurons in the k-th hidden layer such that **W** is *J* × *H*_1_ dimensions (i.e., the number of input variables by the number of neurons in the first hidden layer). Next, let **w**_*j*•_, denote the *j*-th row of the weight matrix **W**. We place a grouped “single-effect” shrinkage prior on the input weights

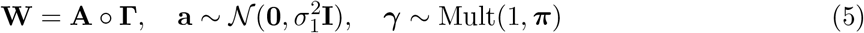

where **Γ** is a matrix that is *H*_1_ copies of the binary vector ***γ***, **a** is a *H*_1_-dimensional row-vector of continuous weights in **A** = [**a**_1•_,…, **a**_*J*•_,], and ○ denotes the Hadamard product between two parameters. Note that this shrinkage prior mimics the sparse assumption of previous neural network architectures in the literature [19, 20], except that the binary indicator variable *γ* is assumed to be multinomial with one trial. Hence, since the *j*-th row of **W** contains the weights connected to the *j*-th column in **X**, when only *γ_j_* = 1, the rest of the input variables are excluded from the model (see proof-of-concept example in Fig. S1 in the Appendix). Together, we refer to the model above as a “single-effect neural network” (SNN). The SNN resembles the SER model in that it assumes that only one input variable has an effect on the response and, thus, posterior summaries of *γ* can be similarly used to compute credible sets.

We now extend the SNN to incorporate multiple effect variables. Analogous to the SuSiE framework, we now consider training on the response variable to be based on an ensemble of single-effect neural networks (ESNN). Probabilistically, the ESNN maybe specified as a summation of *L*-latent nonlinear models of the form

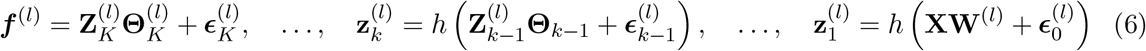

where, in expectation, the response variable is now related to the input data as 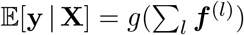 and the sparse prior for the weights of the network are now specified as the following

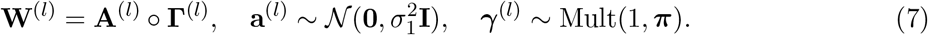

Notice that at the end of training, each *l*-th neural network will also yield an estimated set of input layer weights 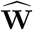 and a corresponding set of inclusion probabilities 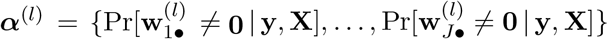 which each assess whether all weights connected to the *j*-th input node are equal to zero. Then, given these *L* posterior summaries, we can compute credible sets 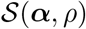 in the same way as SuSiE by defining the overall posterior inclusion probabilities as

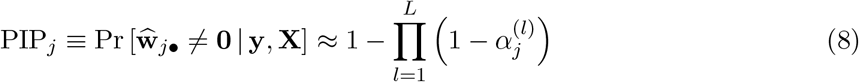

which we use to determine variable significance.

## 4 Posterior Inference via Variational Bayes

As the size of many high-throughput genome-wide sequencing studies continue to grow, both in the number of individuals and the number of genetic variants, it has become less feasible to implement traditional Markov Chain Monte Carlo (MCMC) algorithms for inference. To this end, we use variational inference to approximate the posterior distribution of the weights and hyper-parameters within the ESNN framework. We take the hierarchical model specified in Eqs. (6)-(7) and replace the intractable true posterior distribution over the parameters 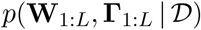 with an approximating family of distributions *q*(**W**_1:*L*_, **Γ**_1:*L*_; ***ϕ***_1:*L*_)—where we use shorthand 1: *L* = 1,…,*L* to represent the *L* models in the ensemble, *ϕ*_1:*L*_ represent the collection of free parameters in the approximations, and 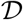 is used to denote the observed data and all relevant hyper-parameters. The basic idea behind the variational inference is to iteratively adjust the free parameters such that they minimize the the difference between the two distributions, which amounts to maximizing the so-called evidence lower bound (ELBO)

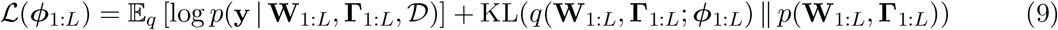

Here, the first term is the expectation of the log-likelihood taken with respect to the variational distribution, and the second term is the Kullback-Leibler divergence which measures the similarity between two distributions. We then use a stochastic gradient descent based method to train models under the ESNN framework. In this work, we choose the variational distributions to factorize across *L* models and for each model we have the following proposals

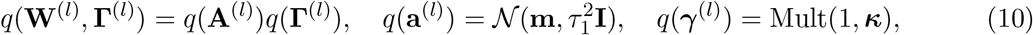

Based on these choices, the gradients of the KL term are available in closed form, while the expectation of the log-likelihood is evaluated using Monte Carlo samples and the local re-parameterization trick (see Appendix for theoretical details and corresponding pseudocode). In a regression task with continuous responses, the log-likelihood term is chosen to be Gaussian and maximizing the lower bound corresponds to minimizing mean square error. In classification tasks for case-control studies, the log-likelihood term is taken to be a binomial distribution which corresponds to minimizing the cross-entropy loss. Since we use gradient descent based method for optimization, the ESNN can be applied for both types of data analyses.

### 4.1 Iterative Bayesian Stepwise Selection

Similar to the SuSiE framework, the ESNN model also uses an iterative Bayesian stepwise selection (IBSS) procedure where it trains *L* models by first fitting one model with a coordinate ascent algorithm and then regressing out that model to compute residuals for training next model. By doing so, we can generate credible sets [9]. It is worth noting that, when the model is uncertain about which variables to choose (e.g., when there are no significant effect variables), ***α*** will become diffuse such that 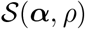 will contain many variables that are not correlated. Under these scenarios, it makes sense to ignore those sets. Previous work have outlined the concept of “purity” as the smallest absolute correlation between all pairs of variables within a credible set which can be used as a criteria for filtering out nonsensical results [9]. This same strategy is not particularly useful on its own for the ESNN framework. An intuitive explanation for this is because since the optimizing objective for neural networks is non-convex, training algorithms can get stuck in local optima where the estimated variational parameters *ϕ* are not optimal. In the scenario where the model is unable to find correct effect variable, regressing out *ϕ* will only introduce noise during training. Therefore, we take an extra approach where we also check to ensure that a trained model is informative before computing the residuals. One simple way to do this is by monitoring whether the likelihood is larger with the *l*-th model trained versus it be excluded from consideration. More specifically, the criteria to include the *l*-th model can be expressed via the (approximate) likelihood ratio

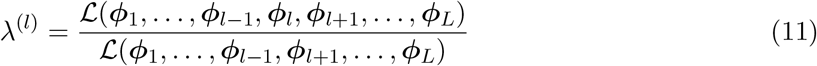

where we keep models that satisfy λ^(*l*)^ > 1. Note that we only regress out variables on continuous data as this is the scenario where it is meaningful to compute the residuals. For the binary classification case, we simply fix the trained models and add up the logits if the criteria is satisfied.

## 5 Results

In this section, we first examine the utility of the ESNN model in simulations motivated by fine-mapping applications for continuous and binary traits in genome-wide association studies. We also apply our method to real world GWA datasets from the Wellcome Trust Case Control Consortium (WTCCC) and the Wellcome Trust Centre of Human Genetics.

### 5.1 Simulations with Continuous Phenotypes

In order to evaluate the performance of our model on continuous traits, we simulate data using real genotypes from chromosome 1 of *N* = 5000 randomly sampled individuals of self-identified European ancestry in the UK Biobank. After quality control [16], this dataset had 36,518 SNPs. To simulate fine-mapping applications, we used the NCBI’s Reference Sequence (RefSeq) database in the UCSC Genome Browser [21] to annotate SNPs to genes. Here, we randomly sampled 200 genes on this chromosome where the annotations included both SNPs located within the gene boundary and SNPs that fall within a ±500 kilobase (kb) window of the boundary to also include regulatory elements.

In this study, each gene is considered to be its own dataset with its own complex correlation structure (see Fig. S3) and unique number of SNPs (ranging from *J* = 50 to 417 variants) encoded as {0,1,2} copies of a reference allele where 0 and 2 represent “homozygotes” and 1 represents “heterozygotes”. For each dataset, we assign 5 effect SNPs and use the following generative model

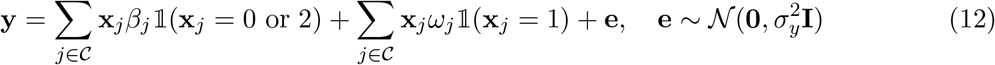

where 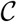 represents the set of causal SNPs and 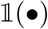 is an indicator function. Here, *β* and *ω* are different effect sizes for heterozygotes and homozygotes, respectively. Both variables are randomly sampled from standard normal distribution and rescaled according to their frequencies. The error term **e** is also assumed to be normally distributed and is rescaled during the simulation such that the causal SNPs explain a certain proportion of the variance in the synthetic trait (i.e., the narrow-sense heritability, *h*^2^). We consider different scenarios where *h*^2^ = {0.05, 0.1, 0.4}.

We compare our method with SuSiE (fit under its default parameter settings). We set *L* = 10 for both approaches. For the ESNN we used a simple sparse architecture with 5 hidden neurons and tanh activation functions. Here, we set the maximum number of epochs to be 30; the hyper-parameter ***π*** for the indicator ***γ*** is chosen from a uniform distribution; we fix 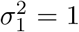 for all *L* models; and, during training, we take 100 Monte Carlo samples to evaluate the log-likelihood. Finally, we used an Adam optimizer with a learning rate of 0.005 and a decay rate of 0.995 after every epoch, and we used an early stopping rule if the likelihood on validation data stopped increasing (based on 85/15 training/validation splits).

To assess the performance, we consider three different metrics. First, we begin by assessing the probability that each method create a credible set containing at least 1 effect SNP (first row Fig. 1(a)). Ideally, a 95% level credible set should have at least 95% coverage. When heritability is high (e.g., *h*^2^ = 0.4), signals are easier to detect, and both ESNN and SuSiE achieve the appropriate coverage. However, for lowly heritable traits (e.g., *h*^2^ = 0.1 and 0.05), the coverage of SuSiE drops while the coverage of ESNN remains. The second metric we check is the average number of effect variables included in all credible sets (first row Fig. 1(b)). In practice, each method can report multiple credible sets. Therefore, this metric essentially helps evaluate the empirical power of ESNN and SuSiE. In these simulations, out method is consistently better regardless of trait heritability. For the final metric, we assess the ability of ESNN and SuSiE to accurately prioritize causal variants according to the PIPs that each method provides. Here, we use receiver operating characteristic (ROC) and precision-recall curves to compare their ability to rank true positives over false positives (first row of Figs. 2 and S5). As *h*^2^ decreases, accuracy of the PIPs for both method decrease but our method is relatively better powered for all scenarios. Importantly, the PIPs from ESNN and SuSiE are calibrated similarly (Fig. S4) [9].

**Figure 1.**
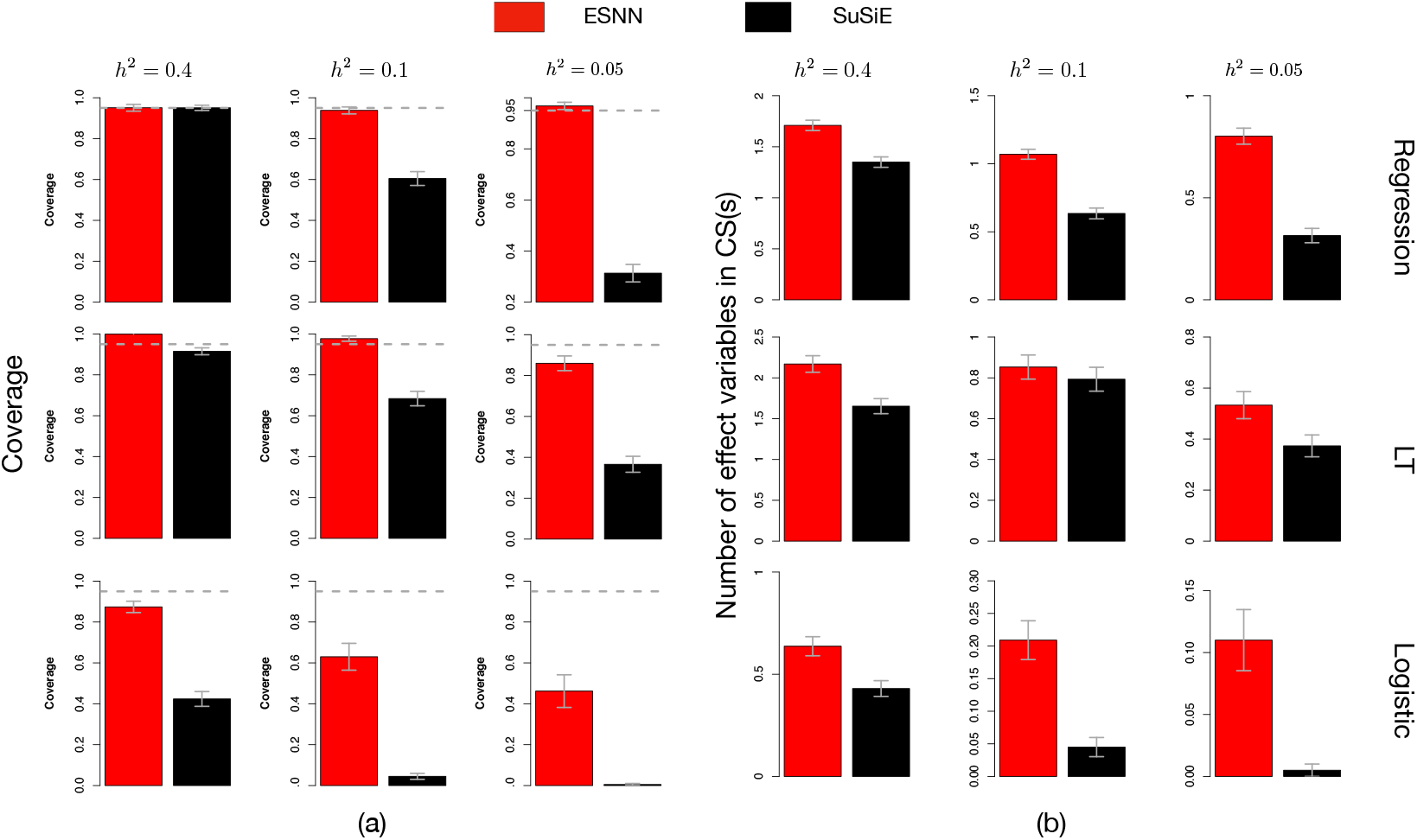
Panel (a) shows comparisons of coverage for ESNN and SuSiE in simulation studies under different levels of heritability. Panel (b) shows the average number of effect variables included in all credible sets for each simulation replicate. Results are based on 200 data replicates.

**Figure 2.**
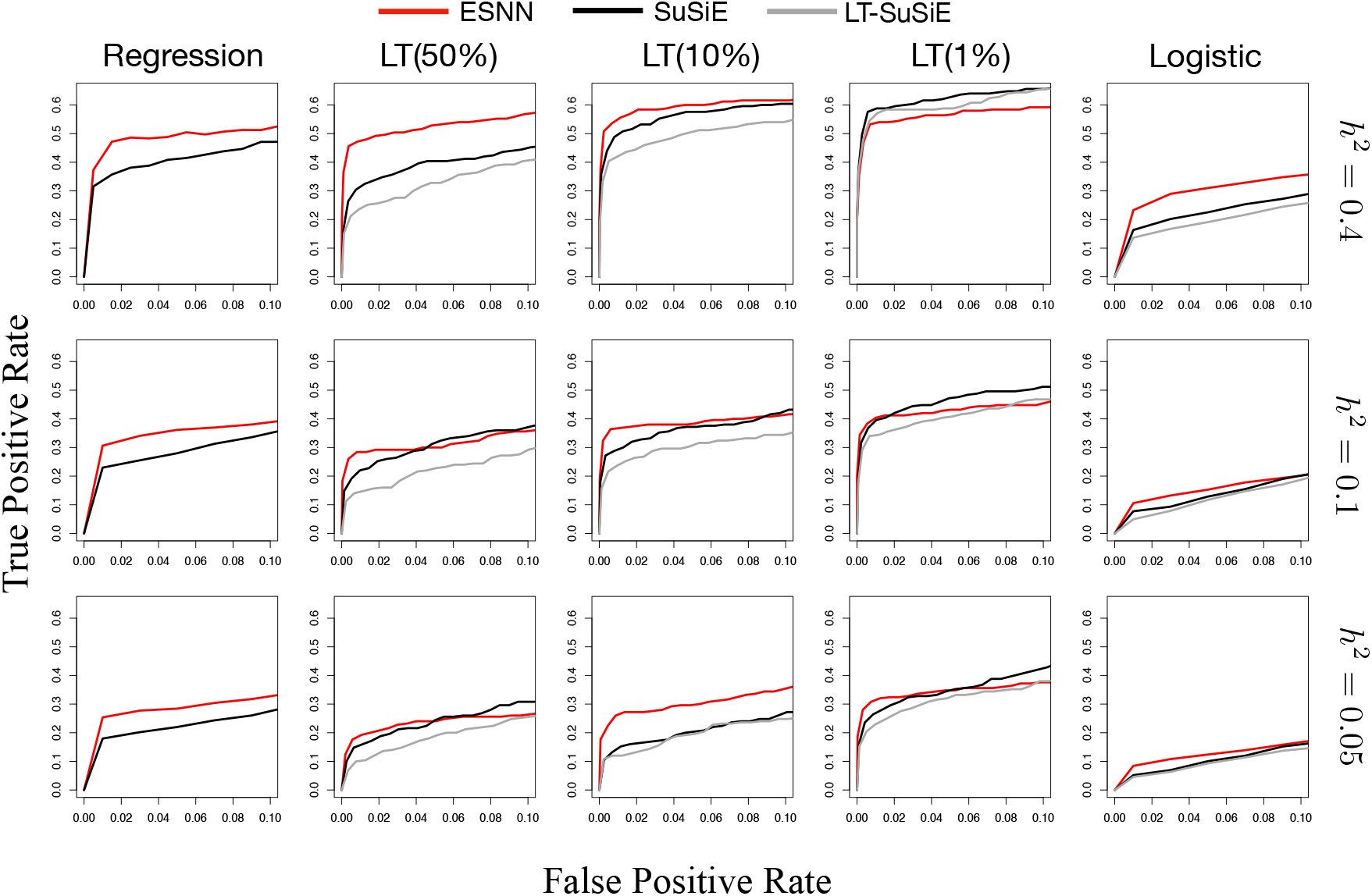
Receiver Operating Characteristic (ROC) curves for simulation studies of different scenarios. Results are based on 200 data replicates.

### 5.2 Simulations with Binary Phenotypes

We now assess the performance of ESNN on binary traits (e.g., case-control studies). We consider two generative models for the class labels: (1) logistic regression and (2) a liability threshold (LT) model [22, 23, 24]. In the former, we simply use the genotypes from chromosome 1 of the *N* = 5000 randomly sampled individuals from the UK Biobank to assume that

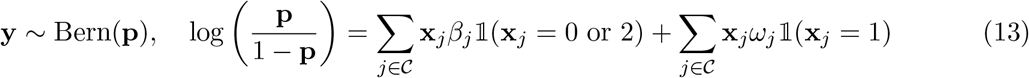

where, in addition to previous notation, the binary traits follow a Bernoulli distribution with probability **p**. In the latter simulation model, we take into account disease prevalence and ascertainment bias which can occur in case-control studies. Here, we adopt the LT model which assumes an latent liability 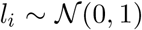 for each observation. With some known prevalence *k*, one can determine a threshold *t* = Φ^-1^(*k*) using the quantile function of normal distribution such that an individual is a case *y_i_* = 1 if *l_i_* > *t*. To simulate data under the LT model, we first generate one million individuals each with *J* = 200 SNPs (with minor allele frequency uniformly sampled between 0.05 and 0.5). Next, we select 5 causal SNPs and generate continuous liabilities with a controlled heritability *h*^2^ = {0.05, 0.1, 0.4} using a model similar to Eq. (12). Then we consider a prevalence *k* ∈ {50%, 10%, 1%} and define cases-controls labels for each of the million individuals. Finally, we subsample 2500 cases and 2500 controls for the analysis.

The SuSiE framework was originally designed for continuous traits, so we consider two adaptations of the model for the binary data. In the first, we simply treat the class labels as continuous and run the model as is. In the second, which we refer to as LT-SuSiE, we use an MCMC to estimate continuous liability scores as phenotypes [25, 24, 26]. Here, we use all the same parameter setting as in the regression simulation study, except that we set the learning rate for ESNN to be 0.01.

Similarly, we compared powers of two methods using coverage (Fig. 1(a)), the number of effect variables included in all credible sets per dataset (Fig. 1(b)), ROC curves (Fig. 2), and precision-recall curves (Fig. S5). Overall, performances follow a similar trend to the regression simulations such that ESNN consistently outperforms SuSiE. When disease prevalence is very low (e.g., k = 1%), cases are assumed to come from “tail” of the distribution. In this scenario, statistical models are generally better powered [27]. As the prevalence k becomes greater, such that the liability threshold moves from the tails to the center of the distribution, it will become harder for a classifier to distinguish cases from controls. This also results in lower power for variable selection. Notably, even in these cases, our method remains robust.

### 5.3 Fine-Mapping in Heterogenous Stock of Mice

We apply ESNN and SuSiE to two continuous traits: high-density and low-density lipoprotein (HDL and LDL, respectively) in a heterogeneous stock of mice dataset from the Wellcome Trust Centre for Human Genetics [28]. This dataset contains *J* = 10,346 SNPs with *N* = 1594 samples for HDL and *N* = 1637 samples for LDL. To run both methods, we simply partition the whole genome into 21 windows where each window contains 500 SNPs. By doing so, we fine-map SNPs in annotated genes as well as SNPs in intergenic regions. We used the same hyper-parameter settings as in the regression simulations for both methods.

For HDL and LDL, ESNN finds 41 and 19 credible sets while SuSiE finds 62 and 26 credible sets, respectively. Our method finding less credible sets is potentially could due to the criteria that we only include an SNN model if it increases the likelihood. This criteria demonstrated to ensure that a credible set generated by ESNN would have high coverage in simulations (Fig. 1). There were 12 SNPs that were included in the credible sets of both methods for HDL and 5 for LDL. This potentially means that these SNPs contributed additive effects to the phenotypic variation. SNPs that are only identified by ESNN probably contribute nonlinear effects (e.g., dominance). We highlighted one region for each trait in Fig. S6. One SNP found by both methods, *rs3090325* in LDL (Fig. S6(a)), can be mapped to the *Smarca2* gene which has been found to be associated with cholesterol regulation [29]. In HDL, SNP *gnf04.147.942* can be mapped to the *Panel* gene which regulates pancreatic activity and has been shown to be linked with HDL [30]. Furthermore, SNPs such as *rs13483562* (which is only found by ESNN in the LDL analysis), can be mapped to the *Aldh1a7* gene which also has been demonstrated to affect related traits such as lipid, cholesterol level, and obesity in mice [31, 32].

### 5.4 Fine-Mapping in the WTCCC 1 Study

We next apply ESNN and SuSiE to two binary traits: type 1 diabetes (T1D) and type 2 diabetes (T2D) from the Wellcome Trust Case Control Consortium (WTCCC) 1 study [33]. This dataset has *N* = 1963 cases and *N* = 2938 controls for T1D, *N* = 1924 cases and *N* = 2938 controls for T2D, along with *J* = 458,868 genotyped SNPs for each individual. Similarly, we run ESNN and SuSiE with a window size of 500 SNPs and used the same model settings as in the binary simulations.

ESNN identifies 32 and 19 credible sets for T1D and T2D, respectively, whereas SuSiE finds and 67 and 30 sets for each trait. There are 5 SNPs that are found by both methods for T1D, but none for T2D. This is likely due to the fact that SuSiE was not originally developed for binary traits and also due to the potential role of nonlinear genetic architecture. We highlight two interesting results in Fig. 3 where we plot the PIPs of SNPs computed by ESNN and SuSiE. In panel (a), we show a window near the *HLA* region on chromosome 6 which has been well studied in the literature and found to be associated with the T1D [34, 35, 36, 37]. One of the two SNPs found only by ESNN, *rs3129051*, is located upstream (within 50kb) of the *HLA-G* gene which is a well-known gene that is related to T1D. The other SNP, *rs16894900*, is located between *MAS1L* (within 50kb downstream) and *UBD* (within 50kb upstream), both of which have been shown to be related to T1D [37]. In panel (b), we highlight the region around *NOS1AP* on chromosome 1. This gene has been found to be linked with T2D in several studies [38, 39, 40]. Our method identified 2 SNPs in this region, but SuSiE reports none. It has been suggested that this region may not play a dominant role in susceptibility to T2D, but a minor effect may exist [38]. Similar to SuSiE, these conclusion were previously made use linear models. We hypothesis this region may contribute to T2D nonlinearly and thus the traditional hypothesis testing methods will have missed this signal.

**Figure 3.**
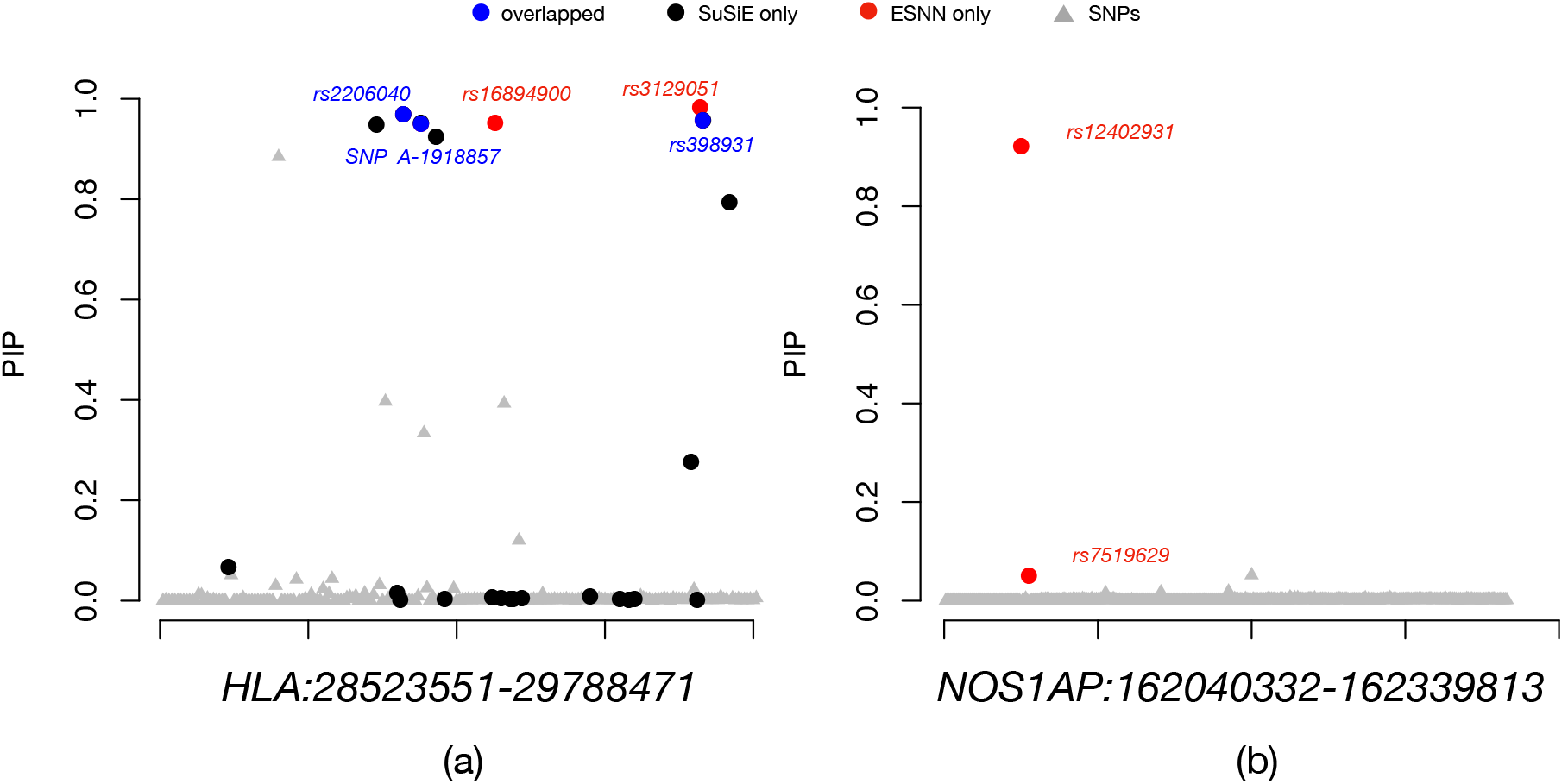
Posterior inclusion probabilities (PIP) of ESNN and SuSiE in the WTCCC analysis. (a) Highlighted region for type 1 diabetes (T1D). Significant SNPs found only by ESNN (included in the credible sets), only by SuSiE, and by both methods are color coded in red, black, and blue, respectively. (b) Highlighted region for type 2 diabetes (T2D).

## 6 Discussion

In this paper, we present the ensemble of single-effect neural network (ESNN) which generalizes the sum of single-effects regression framework by accounting for nonlinear genetic architecture and extending to noncontinuous phenotypes. The ESNN approach provides posterior inclusion probabilities and credible sets that can guide variable selection. While we focus on genetic fine-mapping, this method is also applicable to other fields especially when data are correlated and sparse. We provide a variational algorithm with several relaxation techniques that enables scalable inference. We show that ESNN can effectively increase power for variable selection using simulations. We applied ESNN to two real world genetic datasets and demonstrated its ability to make discoveries that are biologically meaningful. There are a few limitations to the current ESNN framework. Similar to most deep learning models, our method requires large sample sizes for training and requires hyper-parameter fine-tuning. For high-dimensional settings, we currently run the method by splitting the whole dataset into small windows so that the training algorithm can quickly converge. However, this may ignore some long-range interactions. Therefore, a focus for future work will be extending the current model with more complex network architectures.

## Software Availability

Source code and tutorials for implementing the “ensemble of single-effect neural networks” (ESNN) framework are publicly available online at https://github.com/ramachandran-lab/ESNN.

## Data Availability

The heterogenous stock of mice dataset from the Wellcome Trust Centre for Human Genetics can be found at http://mtweb.cs.ucl.ac.uk/mus/www/mouse/index.shtml. Data from the UK Biobank Resource (https://www.ukbiobank.ac.uk) was made available under Application Number 22419. This study also makes use of data generated by the Wellcome Trust Case Control Consortium (WTCCC). A full list of the investigators who contributed to the generation of the data is available from www.wtccc.org.uk. Funding for the WTCCC project was provided by the Wellcome Trust under award 076113, 085475, and 090355.

## Acknowledgements

SR is supported by US National Institutes of Health (NIH) grant R01 GM118652, and National Science Foundation (NSF) CAREER award DBI1452622. LC is supported by a David & Lucile Packard Fellowship for Science and Engineering. Any opinions, findings, and conclusions or recommendations expressed in this material are those of the author(s) and do not necessarily reflect the views of any of the funders.

## Appendix

### 1 Minor Details on the Variational Algorithm

To find the expectation of the log-likelihood during posterior inference, we use Monte Carlo samples and a local re-parameterization trick to compute gradients. More specifically, when assuming Gaussian distributions for the variational approximating families

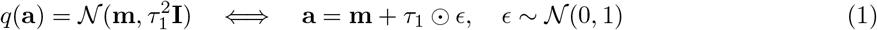

This technique has been shown to successfully reduce the variance of gradients [1] and stabilizes the training process. Next, we assume that the indicator variables ***γ***^(*l*)^ are sampled from a categorical distribution. We adopt a continuous relaxation technique for re-parameterizing these variables by sampling them from a Gumbel-Softmax distribution which is specified as the following [2, 3],

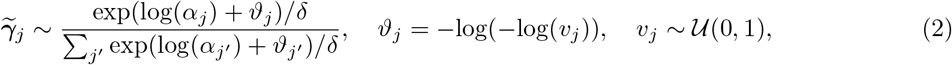

where 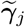 are the approximate samples for ***γ***, *τ* is a temperature parameter, and *v_j_* uniformly sampled random variable. As *τ* → 0, samples 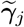 will become closer to the desired vector where only one entry is one and the rest are zeros. In our experiments, we choose *τ* > 0.1 for numerical stability.

The convergence of the inclusion probabilities ***α***_*j*_ is also important for our model as it directly influences the performance of variable selection. Importantly, ***α***_*j*_ appears very early in the computational pipeline since they are defined for the weights in first hidden layer. As a result, the gradients for ***α*** can be very small and hinder convergence during training. This problem is commonly known as “vanishing gradients” [4]. For our work, we found that simply scaling up the learning rate when updating ***α***_*j*_ works well in practice. Note that the Kullback-Leibler (KL) divergence term in the approximate likelihood can be decomposed as the following

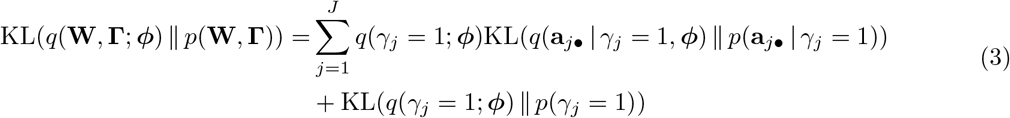

where the KL divergence for the *J* × *H*_1_ weights **W** = **A** ○ **Γ** is between two normal distributions with **A** = [**a**_1•_,…, **a**_*J*•_] and **a**_*J*•_, being an *H*_1_-dimensional row-vector; while the KL divergence for the indicator variables, where ***Γ*** is a matrix that is *H*_1_ copies of the *J*-dimensional binary vector **γ**, is taken between two discrete multinomial distributions. Importantly, these terms have closed-form solutions with which gradients can be computed.

### 2 Pseudocode for ESNN

**Algorithm 1.**
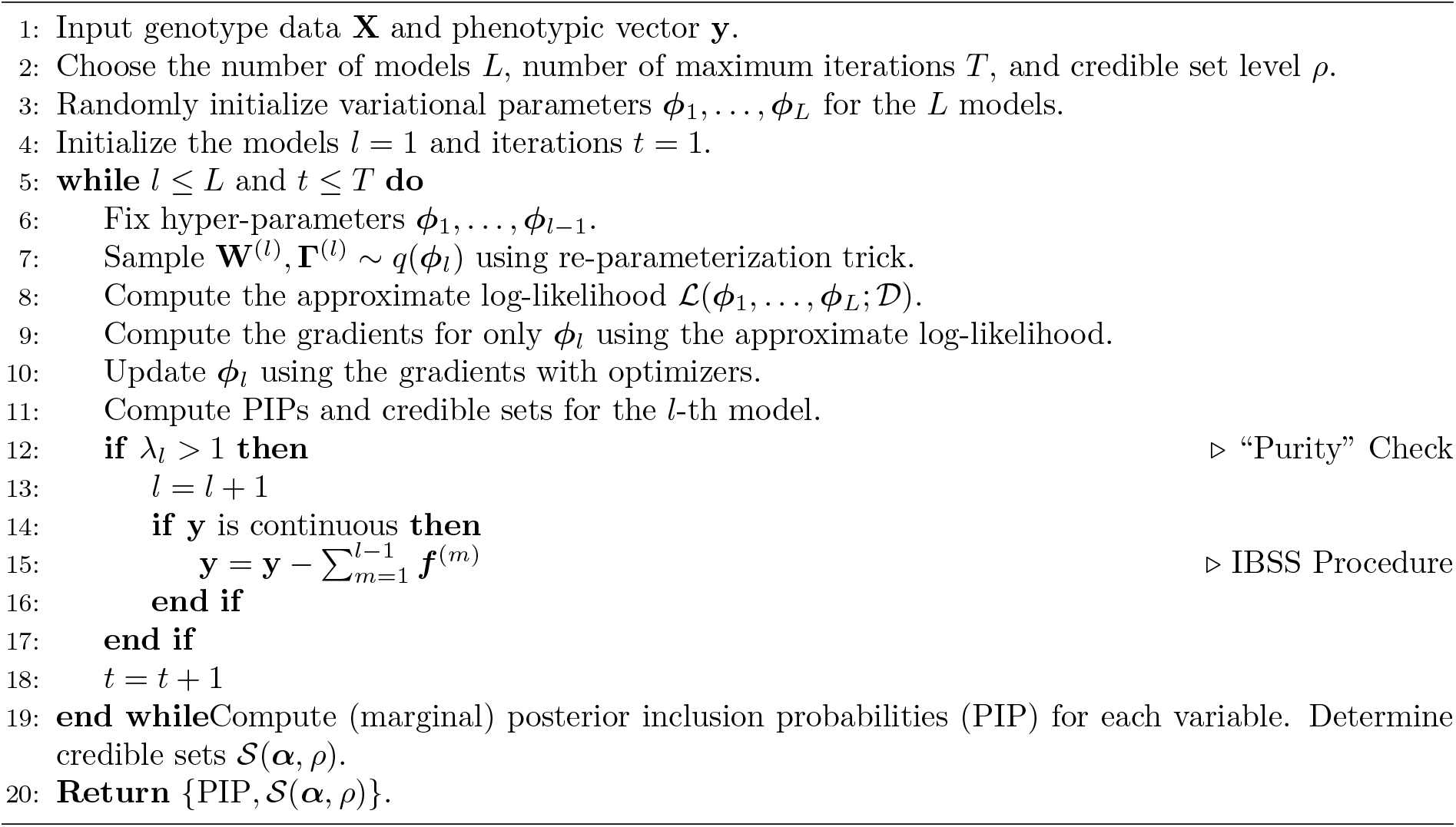
Training Algorithm for the ESNN Framework

### 3 Supplementary Figures

**Figure S1.**
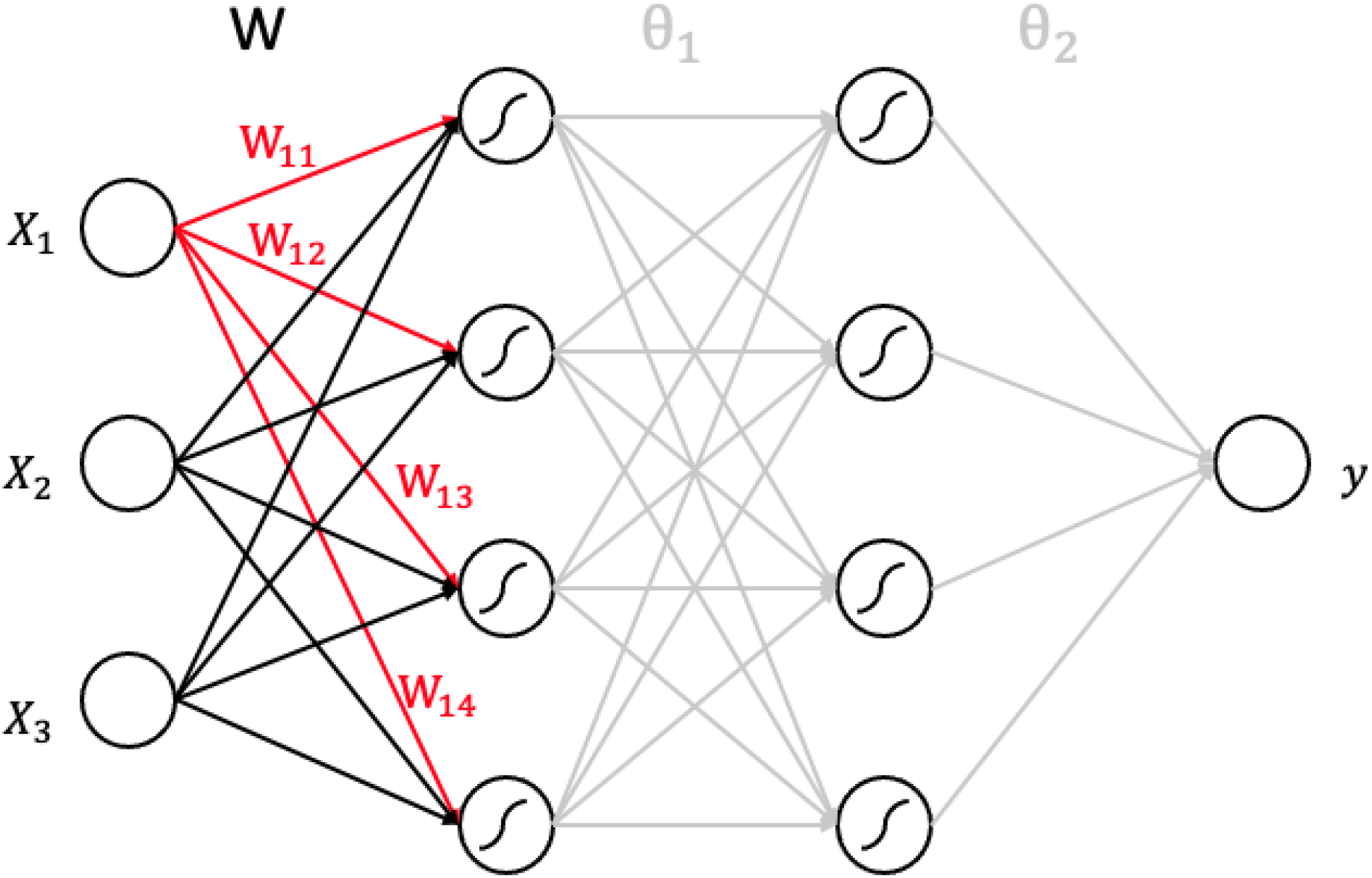
An example of a single-effect neural network (SNN) with only the first input variable having an effect on the outcome.

**Figure S2.**
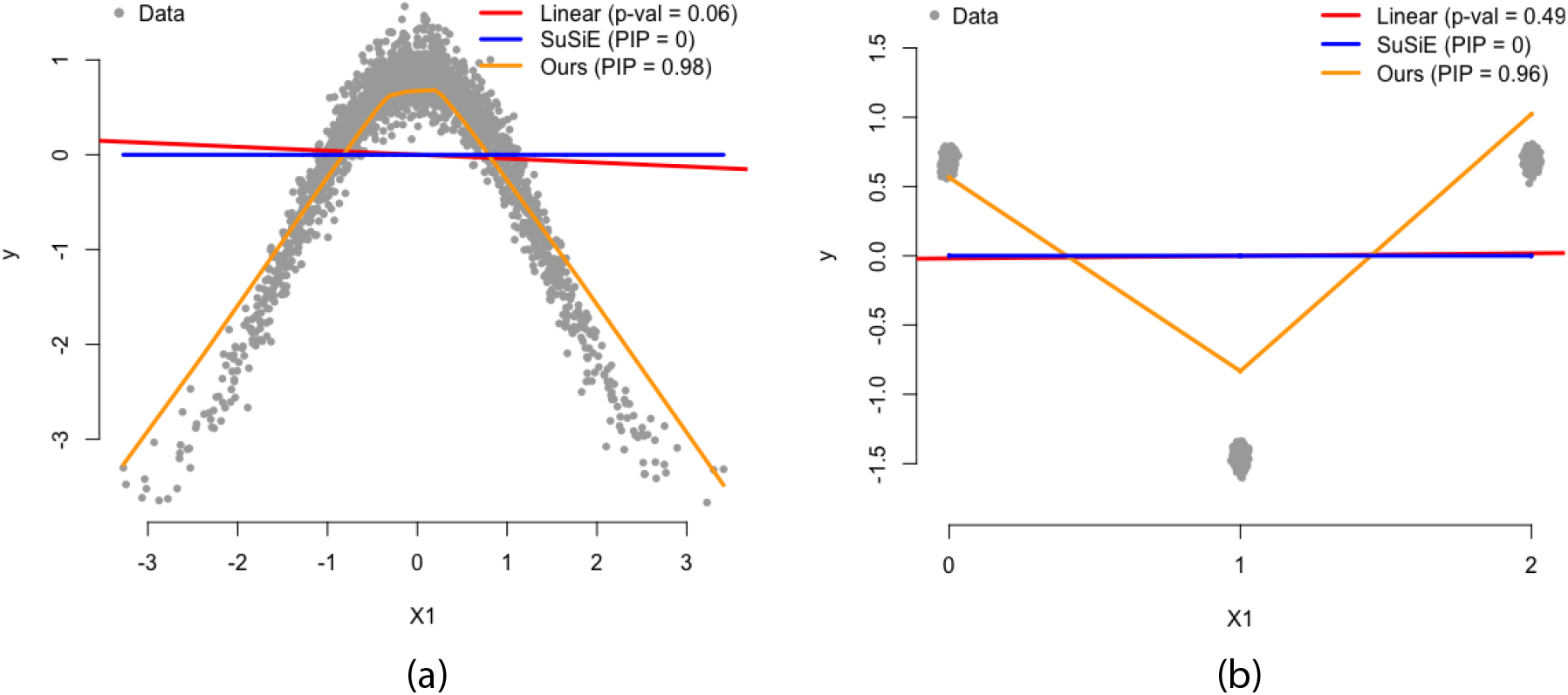
Toy example demonstrating importance of accounting for nonlinearity when performing variable selection. To demonstrate that linear models lose power when non-additive variation exists, we generate two simulated datasets. **(a)** In the first case, we simulate 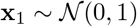 and then generate responses under **y** = cos(**X**_1_) + **e** where 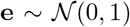. The real data are plotted in grey points. We next run a univariate linear model (red line) and SuSiE (blue line) [5] on this dataset. Here, we perform variable selection by ranking the resulting p-values and posterior inclusion probabilities (PIP) for the respective approaches. These results show that neither method selects **x**_1_ as being significant. We then run the ESNN (orange line) on these data and it successfully captures the signal. **(b)** In the second simulation example, we mimic a genome-wide association (GWA) study. Here, we use single nucleotide polymorphisms (SNPs) with values taking on {0, 1, 2} based on copies of a reference allele where 0 and 2 represent “homozygotes” and 1 represents “heterozygotes”. We then simulate the phenotype **y** by assuming the heterozygote has a significant effect. Similar relationships have been shown in the literature [6,7]. Similar to the first simulation case, linear methods fail to capture the causal effect while the ESNN is robust to the non-additive architecture. These two toy examples illustrate the importance of accounting for nonlinearity in variable selection methods.

**Figure S3.**
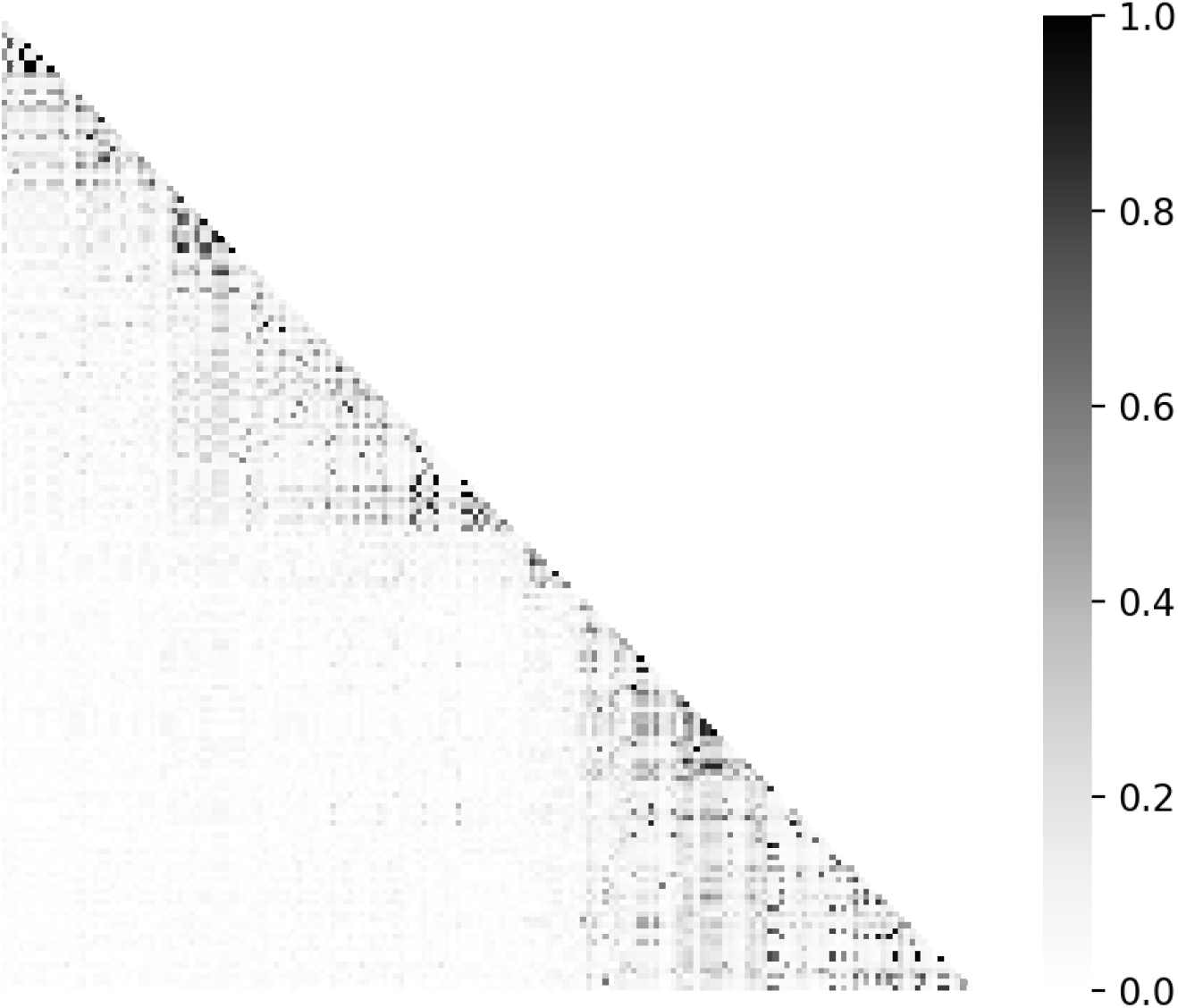
An example of absolute correlation matrix for genotype data.

**Figure S4.**
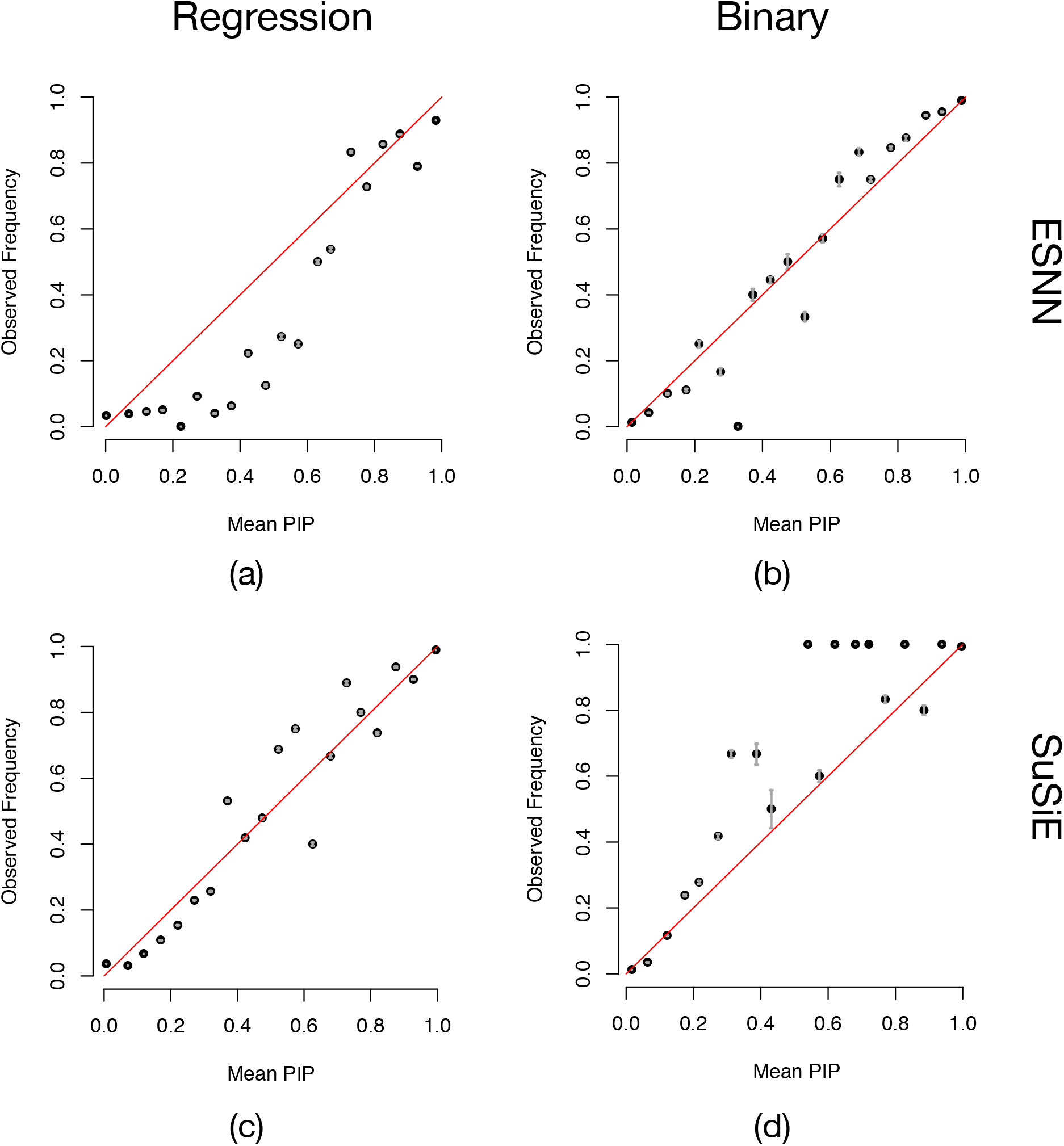
Assessments of posterior inclusion probability (PIP) calibration for ESNN and SuSiE. This experiment follows largely from previous work [5]. Here, SNPs are grouped into bins according to their reported PIPs (using 20 equally spaced bins, from 0 to 1).The plots show the average PIP for each bin against the proportion of causal SNPs or SNP-sets in that bin. A well calibrated method should produce points near the x-axis = y-axis line (i.e., the diagonal red lines). Gray error bars show ±2 standard errors. Panel **(a, b)** shows the comparison of ESNN and panels **(c, d)** shows the comparison of SuSiE for continuous and binary traits, respectively.

**Figure S5.**
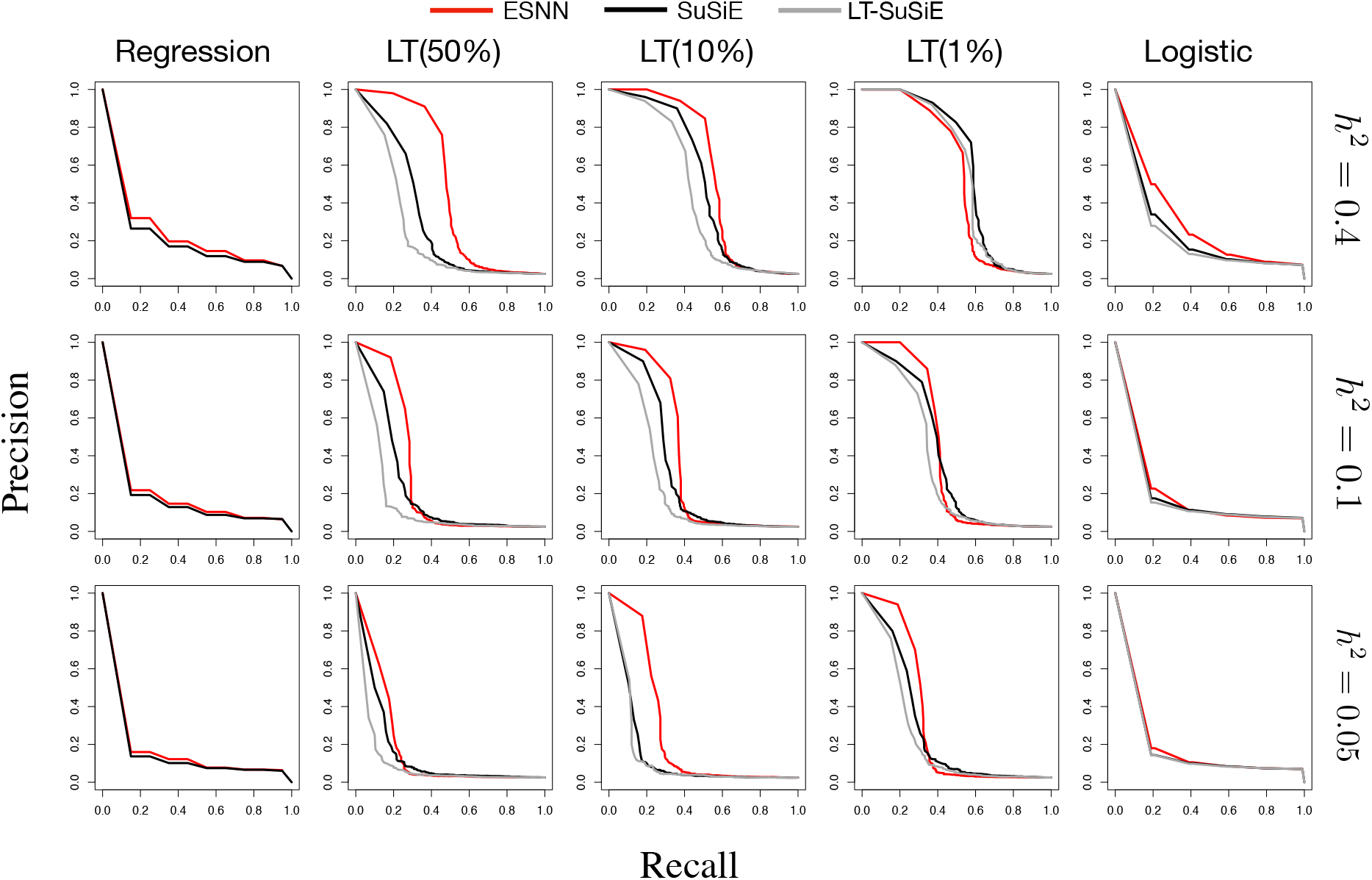
Precision and recall curves for simulation studies of different scenarios. Results are based on 200 data replicates.

**Figure S6.**
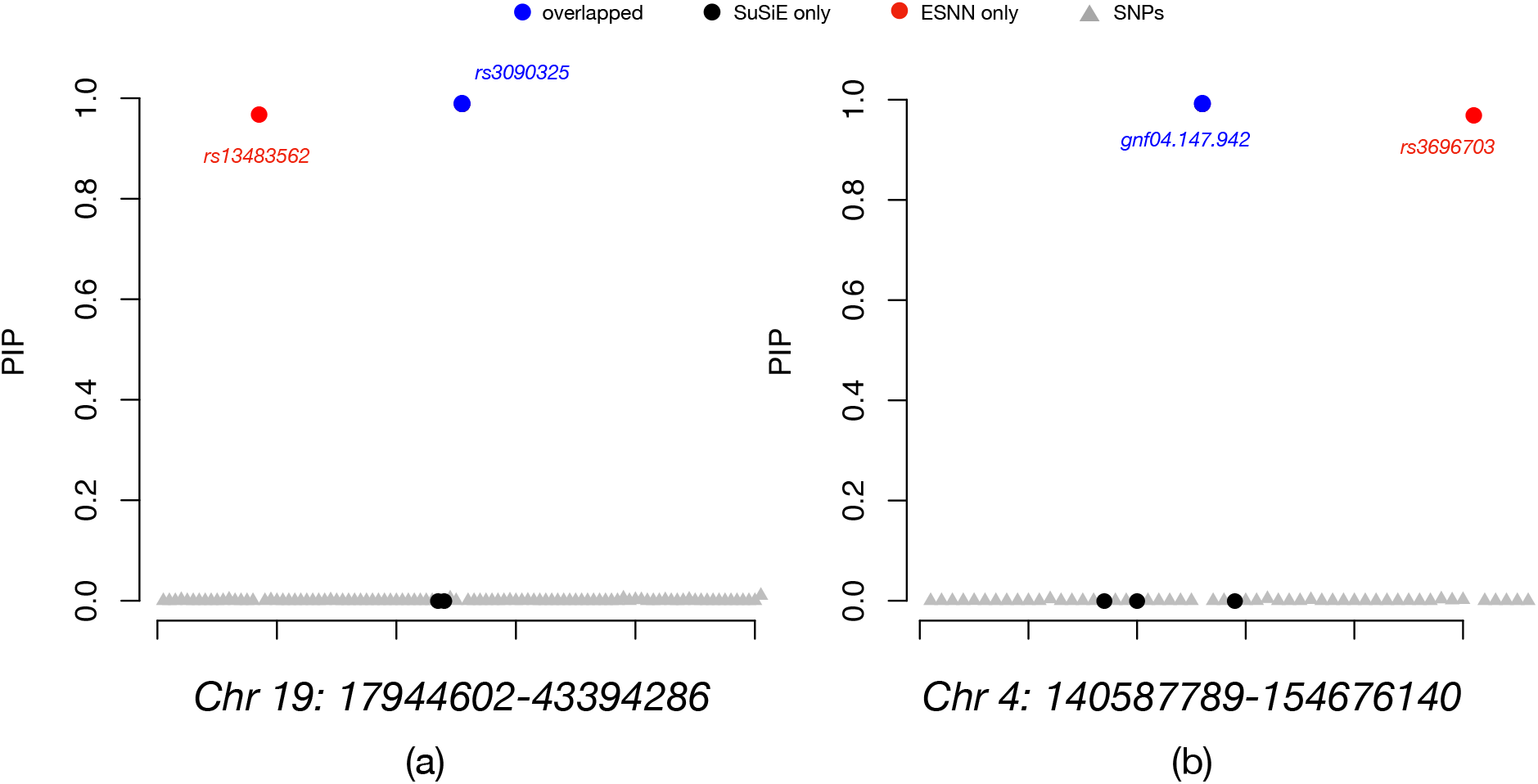
Posterior inclusion probabilities (PIP) of ESNN and SuSiE for the heterogenous stock of mice dataset from the Wellcome Trust Centre for Human Genetics [8]. **(a)** Highlighted region for low-density lipoprotein (LDL). Significant SNPs found only by ESNN (included in the credible sets), only by SuSiE, and by both methods are color coded in red, black, and blue, respectively. **(b)** Highlighted region for high-density lipoprotein (HDL).

